# Mutations affecting the N-terminal domains of SHANK3 point to different pathomechanisms in neurodevelopmental disorders

**DOI:** 10.1101/2021.04.13.439653

**Authors:** Daniel Woike, Emily Wang, Debora Tibbe, Fatemeh Hassani Nia, Maria Kibæk, Martin J. Larsen, Christina R. Fagerberg, Igor Barsukov, Hans-Jürgen Kreienkamp

## Abstract

Shank proteins are major scaffolds of the postsynaptic density of excitatory synapses. Mutations in *SHANK* genes are associated with autism and intellectual disability. The relevance of missense mutations for these pathologies is unclear. Several missense mutations in *SHANK3* affect the N-terminal region, consisting of the Shank/ProSAP N-terminal (SPN) domain and a set of Ankyrin (Ank) repeats. Here we identify a novel *SHANK3* missense mutation (p.L270M) in the Ankyrin repeats in patients with an ADHD-like phenotype. We functionally analysed this and a series of other mutations, using biochemical and biophysical techniques. We observe two major effects: (i) a loss of binding to δ-catenin (e.g. in the p.L270M variant), and (ii) interference with the intramolecular interaction between N-terminal SPN domain and the Ank repeats. This also interferes with binding to the α-subunit of the calcium-/calmodulin dependent kinase II (αCaMKII), and appears to be associated with a more severe neurodevelopmental pathology.

## Introduction

Neurodevelopmental disorders are frequently caused by mutations in genes coding for proteins which contribute to the formation and function of synapses. This has led to the term “synaptopathies” (1, 2). Genetic alterations affect both pre- and postsynaptic components. One prominent family of genes, which has early on been implicated in the etiology of autism, intellectual disability and schizophrenia is the *SHANK* family of genes, coding for SHANK 1-3 (2-4). Loss of one copy of the *SHANK3* gene is observed in 22q13 deletion syndrome/Phelan McDermid syndrome; in addition to loss-of-function mutations, also numerous missense mutations have been described (4-6). Shank proteins are multidomain scaffold proteins of the postsynaptic density of excitatory, glutamatergic synapses (7). They connect, through numerous protein interactions, the glutamate receptors of the synapse with the F-actin based cytoskeleton of the dendritic spine (8). In detail, the central PDZ domain of Shank proteins binds to SAPAP/GKAP family members, which links Shank to the PSD-95/NMDA receptor complex (9, 10). Multimerization of the C-terminal SAM domains of Shank2 and Shank3 is believed to be essential for PSD formation (11, 12). A long proline rich region between these two domains provides anchoring points for cytoskeleton-associated proteins such as IRSp53, cortactin, Abi1, as well as Homer, a scaffold for metabotropic glutamate receptors (10, 13-15). Somewhat surprisingly, few (if any) of the many missense mutations in *SHANK* genes found in patients affect these “core” motifs and domains of Shank proteins. Instead, missense mutations in the C-terminal half of Shank proteins lead to alterations outside of known interaction motifs, making it difficult to assess the functional relevance of these mutations (16).

The N-terminal region of Shank3 has emerged as a hotspot for missense mutations (6). This region includes the Shank/ProSAP N-terminal domain (SPN) and a set of seven ankyrin repeats (Ank). Recent structural work has shown that the SPN domain is folded as a ubiquitin like (Ubl) domain, similar to Ras association domains, and high affinity binding of active, GTP-bound variants of Ras and Rap to the SPN domain of Shank3 has been demonstrated (17, 18). Mutations R12C and L68P in the SPN domain which were found in autistic patients disrupt G-protein binding (17).

The Ank repeats interact with postsynaptic partners as α-Fodrin and δ-catenin (19, 20), as well as Sharpin (the postsynaptic localization of which has been unclear) (21). In addition, the SPN domain is in close intramolecular contact to the Ank domain, to the effect that the SPN domain interferes with the access of some known interaction partners of the Ank repeats, namely α-Fodrin and Sharpin (17, 22). Furthermore, recent biochemical work demonstrated an interaction of the inactive, non-phosphorylated form of αCaMKII with the loop between SPN and Ank, partially covering the Ras binding site (23). The functional relevance for most of these interactions for the synaptic function of Shank proteins remains largely unclear. More importantly, it remains to be clarified how individual missense mutations in the N-terminal part of Shank3 lead to neurodevelopmental disorders, such as autism and schizophrenia.

Here we perform a functional analysis of missense mutations affecting the N-terminal SPN and Ank domains of Shank3. Mutations are taken from novel patients presenting with attention deficit/hyperactivity disorder (ADHD) and autistic phenotypes, from the ClinVar database and from the published literature. Our experiments identify two major aspects of ASD-associated mutations: interference with the tight association of SPN and Ank domains, and reduced binding to δ-catenin.

## Materials & Methods

### Whole exome sequencing (WES)

DNA was extracted from EDTA-stabilized peripheral blood lymphocytes and subjected to exome capture using NimbleGen SeqCap EZ MedExome (Roche), followed by sequencing on an Illumina NovaSeq platform to a mean coverage of minimum 300x, with 95% of targeted bases covered with a minimum coverage of 30x. Raw reads were aligned using the Burrows-Wheeler Alignment tool v. 0.7.15 and the GATK (Genome Analysis Toolkit) Best Practice pipeline v. 3.8–0 was used for variant calling (24). Annotation and filtering of variants was performed using VarSeq 2.2.0 (Golden Helix). The sequence variant has been confirmed by bidirectional Sanger sequencing according to standard procedures. Written consent for publication of these data has been obtained from all individuals or their legal guardians presented in this study.

### DNA constructs

Bacterial expression constructs coding for His_6_/SUMO-tagged fusion proteins of rat Shank3 (residues 1-348, SPN-Ank) were generated in pET-SUMO (Thermo Fisher Scientific) as described before (17). Constructs coding for N-terminal (residues 1-339, SPN-Ank; residues 75-339, Ank only) rat Shank3 with a C-terminal mRFP-tag were generated in pmRFP-N3 (Clontech). A construct coding for N-terminally GFP-tagged full-length rat Shank3 in the pHAGE vector was obtained from Alex Shcheglovitov (Univ. of Utah, Salt Lake City, UT) (17, 25). For live FRET imaging, rat Shank3 N-terminal constructs (WT, R12C, N52R, L68P and P141A) coding for amino acids 1-339 of Shank3 were used, including an N-terminal GFP and a C-terminal mCherry sequence. Missense mutations were introduced by site-directed mutagenesis using the Quik-Change II site-directed mutagenesis kit (Agilent) and mutagenic oligonucleotides obtained from Sigma-Aldrich. Construct identity was confirmed by Sanger sequencing. A construct coding for HA-tagged HRas G12V was obtained from Georg Rosenberger (UKE Hamburg, Germany). cDNA coding for Rap1a G12V was cloned into the pEGFP-C1 vector (Clontech). An expression vector coding for mouse δ-catenin carrying an N-terminal GFP-tag in pEGFP-C1 was obtained from K. Kosik (Univ. of California, Santa Barbara, CA). Constructs coding for the C-terminal part of α-Fodrin fused to an N-terminal EGFP-tag (19) and full length αCaMKII in a modified pcDNA3 vector coding for an N-terminal T7-tag (26) have been described before.

### Cell culture and transient transfection

Human embryonic kidney (HEK) 293T cells were cultured in Dulbecco’s Modified Eagle Medium supplemented with 10% fetal bovine serum and 1% Penicillin/Streptomycin. A maximum of 20 cell passages were used. Transient transfection of 293T cells was performed using TurboFect Transfection Reagent (Thermo Fisher Scientific) according to the manufacturer’s instructions.

### Cell lysis and immunoprecipitation

Cell lysis was performed using immunoprecipitation (IP) buffer (50 mM Tris pH 8, 120 mM NaCl, 0.5% NP40, 1 mM EDTA). Immunoprecipitation was performed using RFP-trap beads (Chromotek). Precipitates were washed five times in IP buffer; both input and precipitate samples were then processed for SDS-PAGE and Western blotting.

### SDS-PAGE and Western blot

Proteins were denatured in 1x Laemmli sample buffer (63 mM Tris/HCl, pH 6.8; 10% glycerol; 1.5% SDS; 0.1 M Dithiothreitol; and 0.01% bromophenol blue) at 95 °C, separated on 10% SDS-PAGE at 100–180 V and transferred to nitrocellulose membrane in transfer buffer (25 mM Tris; 192 mM glycine; 0.05% SDS; and 20% methanol) at 100 V for 100 min using a MINI PROTEAN II^TM^ system (Bio-Rad). Membranes were blocked with 5% milk powder/TBS-T (Tris buffered saline; 10 mM Tris/HCl pH 8; 150 mM NaCl, with 0.05% Tween 20) and incubated with the indicated primary antibodies overnight at 4 °C followed by HRP-linked secondary antibodies at room temperature for 1 h. After washing the membranes with TBS-T, chemiluminescence was detected using WesternBright chemoluminescence substrate (Biozym, Hess. Oldendorf, Germany). Membranes were scanned using a ChemiDoc^TM^ MP Imaging System (Bio-Rad) and images were processed and analyzed using Image Lab Software (Bio-Rad).

### Animals

For preparing primary neuronal cultures, brain tissue was isolated from *Rattus norvegicus* embryos. Pregnant rats (Envigo; 4-5 months old) were sacrificed on day E18 of pregnancy using CO_2_ anaesthesia, followed by decapitation. Neurons were prepared from all embryos present, regardless of gender (12-16 embryos). All animal experiments were approved by, and conducted in accordance with, the guidelines of the Animal Welfare Committee of the University Medical Center (Hamburg, Germany) under permission number Org766.

### Hippocampal neuron culture and transfection

Primary hippocampal neurons were isolated from E18 rat embryos. The hippocampal tissue was dissected, and hippocampal neurons were extracted by enzymatic digestion with trypsin, followed by mechanical dissociation. Cells were grown in Neurobasal medium supplemented with 2% B27, 1% Glutamax and 1% Penicillin/Streptomycin. Neurons were transfected on DIV7 using the calcium phosphate method. The complete Neurobasal medium was collected from wells one hour before transfection and replaced with transfection medium (Minimal Essential Medium + Glutamax). 5 μg plasmid DNA was mixed with 10 μL 2.5 M CaCl_2_ and topped up to 100 μL with ddH_2_O. An equal amount of 2x Hepes buffered salt solution (HBS pH 7.05; 274 mM NaCl; 10 mM KCl; 1.4 mM Na_2_HPO_4_; 15 mM D-Glucose; 42 mM Hepes) was added dropwise to the reaction tube under continuous vortexing. The reaction was incubated at room temperature for 30 min and then divided between the wells of the cell culture plate. After a 2 h incubation with the transfection mixture, the cells were washed seven times with 1x Hank’s Balanced Salt Solution (HBSS). After the final wash, the previously collected Neurobasal medium was added back to the cells.

### Immunocytochemistry

Neurons (DIV14; seven days after transfection) were fixed with 4% paraformaldehyde, 4% sucrose in PBS and permeabilized with 0.1% Triton X-100 in PBS for 5 min at room temperature. After blocking (10% horse serum in PBS) for 1 h at room temperature, neurons were incubated with corresponding antibodies overnight followed by 1 h of incubation with Alexa Fluor antibodies. The coverslips were mounted onto glass microscopic slides using ProLong^TM^ Diamond Antifade mounting medium (Thermo Fisher Scientific).

### Live FRET (Förster resonance energy transfer) imaging

293T cells were transfected with FRET constructs and imaged live in Hank’s balanced salt solution (HBSS; without phenol red) on the following day for 1 min The FRET coefficient was calculated point by point in the image by making the following operation between the acceptor channel (mCherry) and the donor channel (GFP):

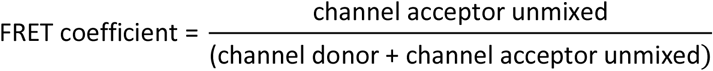

FRET coefficient values range between 0 and 1. To display them as 8-bit images, these values were rescaled ranging between 0 and 255. The value 1 (255) represents maximum FRET efficiency while 0 is equivalent to no detectable FRET.

### Expression and purification of recombinant protein

Shank3 protein (residues 1-348; WT and P141A mutant) as a fusion with a His_6_-SUMO-tag was expressed from pOPINS vectors (OPPF-UK) in the BL21*(DE3) E. coli strain. Cultures were grown in LB media supplemented with antibiotics. Cells were grown at 37 °C to an optical density at 600 nm of 0.6-0.7 and cooled to 18 °C before inducing with 0.5 mM isopropyl-β-D-1-thiogalactopyranoside (IPTG) overnight. Cells were harvested by centrifugation at 12000 × g, resuspended in low imidazole buffer (10 mM imidazole, 500 mM NaCl, 20 mM Na_2_HPO_4_, pH 7.4 with 10% glycerol v/v), frozen and stored at −20 °C. Cells were lysed through treatment with lysozyme (0.2 mg/mL) in the presence of protease inhibitor cocktail and 2 mM dithiothreitol (DTT) (2 mM) for 30 minutes before sonication on a Soniprep 150 Plus Disintegrator (MSE). After centrifugation at 60,000 × g the supernatant was used for purification using nickel-affinity chromatography on a HisTrap HP column (GE Healthcare), using an isocratic elution of low to high imidazole buffer (500 mM imidazole, 500 mM NaCl, 20 mM Na_2_HPO_4_, pH 7.4). Eluted protein was then cleaved using recombinant SUMO protease, followed by overnight dialysis. The His-SUMO-tag was removed by a reverse pass on the HisTrap HP column. Proteins were exchanged into 1 M NaCl, 20 mM Tris, 2 mM DTT pH 7.4.

### Differential scanning fluorimetry (DSF)

Proteins were diluted to 5 uM in varying salt conditions ranging from 0.125 M to 1 M NaCl, 20 mM Tris pH 7.5. Samples were added to wells in a MicroAmp Fast 96 well reaction plate (Applied Biosystems), along with a 1:1000 dilution of Sypro Orange Protein Gel Stain (Invitrogen) and a 1:10 dilution of 200 μM Tris pH 7.5. Thermal stability was then measured on a StepOnePlusTm Real-Time PCR System (Applied Biosystems) over a temperature range of 25 °C to 95 °C. Results were analysed using StepOneTm melting software (Applied Biosystems). Smoothed melting curve data were taken and the normalized mean fluorescence for each sample was plotted to generate a melting transition curve. The transition curve was isolated and a Boltzmann statistical analysis was done to give a Tm for each sample.

### Microscopy

Confocal images were acquired with Leica SP5/SP8 confocal microscopes using a 63x objective. Quantitative analysis for neuron images was performed using ImageJ. The evaluation of the fluorescence resonance energy transfer was performed using Bitplane Imaris software.

### Antibodies

The following primary antibodies were used: mouse anti-GFP (Covance MMS-118P; RRID:AB_291290; WB: 1:3000), mouse anti-HA-7 (Merck H9658; RRID:AB_260092; WB 1:20000) mouse anti-T7-tag (Merck 69522; RRID:AB_11211744; WB 1:10000) and mouse anti-PSD-95 (Thermo Fisher Scientific MA1-046, RRID:AB_2092361; ICC: 1:500); rat anti-RFP (Chromotek 5F8; RRID:AB_2336064; WB: 1:1000); chicken anti-MAP2 (Antibodies-Online; RRID:AB_10786841; ICC: 1:1000). HRP-labeled goat secondary antibodies were from Jackson ImmunoResearch (RRID:AB_2338133; RRID:AB_10015289) and used for WB at 1:2500 dilution. For ICC, Alexa 633 goat anti-mouse IgG (Invitrogen A21050; RRID:AB_2535718) and Alexa 405 goat anti-chk IgG (abcam ab175675; RRID:AB_2819980) were used at 1:1000 dilution.

### Protein stability prediction

Protein stability analysis was performed *in silico* using the PoPMuSiC program (27) available at soft.dezyme.com.

### Evaluation of data

All data are presented as mean ± SD. No statistical method was used to determine sample size. Sample size was based on experience with the relevant type of experiment (17, 20, 22, 26), and with the objective to minimize the number of animals killed. No randomization methods were used. No test for outliers was conducted on the data; no data points were excluded. Statistical significance was determined using Prism8 software (GraphPad, San Diego, CA) and analysed by Student’s t-test or one-way ANOVA with *post hoc* Dunnett’s test.

## Results

### A *SHANK3* missense mutation with an unusual phenotype

We identified a family with several individuals affected by learning difficulties, an oppositional behavioral disorder and ADHD–like behavior. Upon exome sequencing, we identified a heterozygous missense mutation in the *SHANK3* gene (NM_033517.1:c.808C>A, p.Leu270Met). The variant appeared to segregate with the ADHD or ADHD-like symptoms in the family (Fig. 1a). ADHD has so far not been included in the spectrum of *SHANK3*-associated diseases. To clarify the pathogenicity of this variant in *SHANK3* we began to compare the functional effects of this mutation with those of other variants in *SHANK3* which are associated with autism spectrum disorders (ASD), intellectual disability (ID) or other disorders such as epilepsy. To do so, we performed a systematic analysis of this mutation and other patient missense mutations in *SHANK3* affecting the N-terminal region of the Shank3 protein which had not been functionally analyzed before (Fig. 1b). We included the P141A mutation found in an autistic individual (28). In addition, we included all missense mutations found in the ClinVar database which altered residues in the N-terminal SPN and Ank domains, and which were absent from the gnomAD database of 125,748 exome sequences and 15,708 whole-genome sequences (29). One variant (I245T) which was listed in ClinVar was present many times in gnomAD and was therefore excluded.

**Figure 1.**
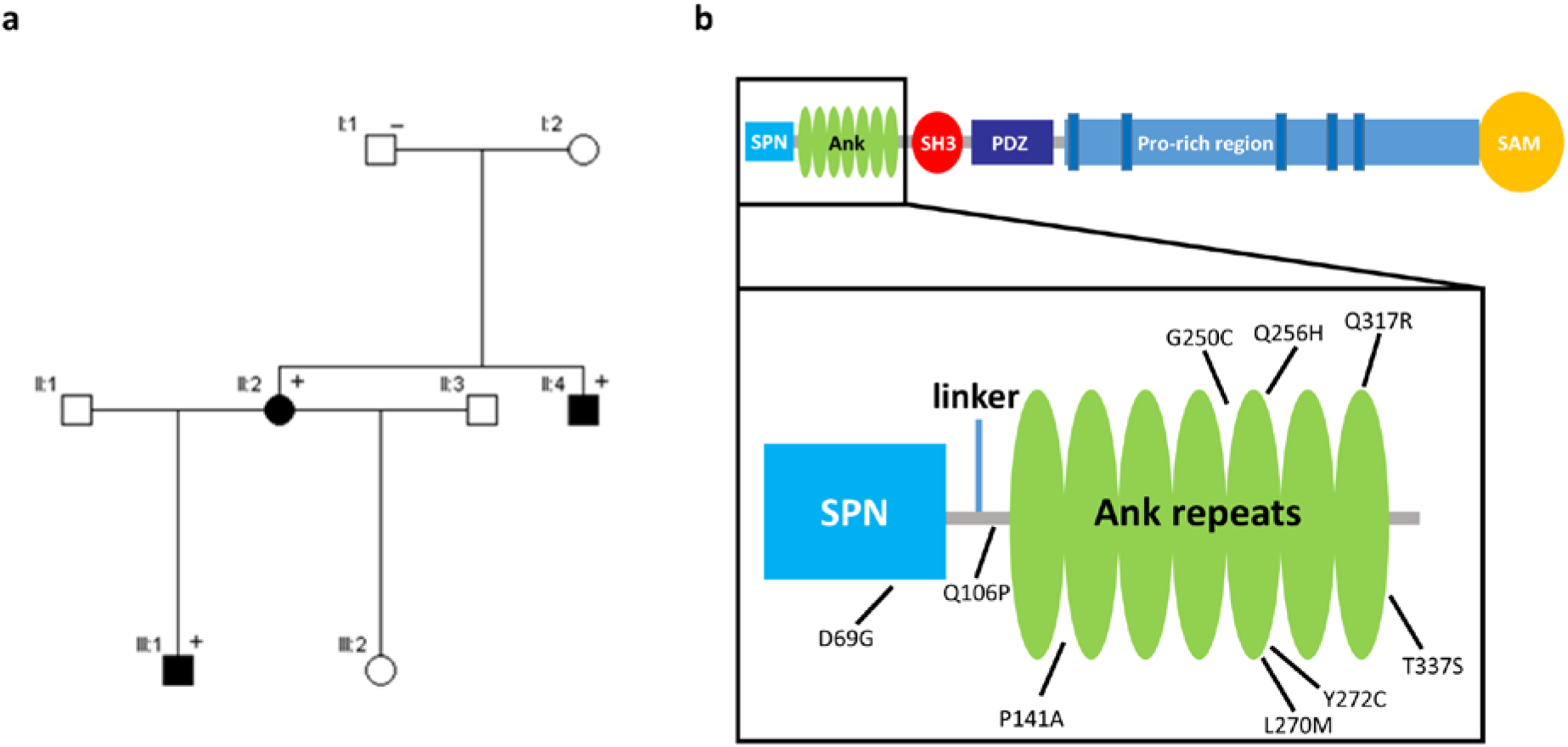
**a**. Pedigree of the family with the p.L270M *SHANK3* missense variant. +/- denotes presence/absence of the variant. **b**. Domain structure of Shank3. The positions of the N-terminal missense mutations examined in this study are indicated.

An initial bioinformatic analysis of the effect of the mutations on protein stability was performed based on the crystal structure of the Shank3 N-terminus (5G4X, (17)). All mutations had a negative impact on thermodynamic stability of the protein, represented as positive ΔΔG values in Table 1. The P141A, G250C and Y272C mutations led to the highest ΔΔG values, whereas T337S had the smallest effect on protein stability.

**Table 1.** Overview of SHANK3 variants analyzed in this study. Solvent accessibility of the residue at this position and free energy changes (ΔΔG values), indicating altered stability, are calculated using PoPMuSiC software.

| Variant, domain | Source | Phenotype | Interpretation | Inheritance | Solvent accessibility (%) | $\Delta\Delta G$ |
| --- | --- | --- | --- | --- | --- | --- |
| D69G SPN | ClinVar | Autism | Unknown significance | *De novo* | 55.77 | 0.64 |
| Q106P linker | ClinVar | Phelan McDermid syndrome* | Unknown significance | Not provided | 50.96 | 1.28 |
| P141A Ank | Boccuto et al., 2013 | Developmental delay, seizures, autism | Pathogenic | *De novo* | 2.21 | 1.33 |
| G250C Ank | ClinVar | Not provided | Likely pathogenic | Not provided | 24.18 | 1.73 |
| Q256H Ank | ClinVar | Not provided | Likely pathogenic | Not provided | 5.98 | 0.87 |
| L270M Ank | This study | Learning difficulties; oppositional behavioral disorder; ADHD-like behavior | Unknown significance | Inherited | 3.01 | 1.11 |
| Y272C Ank | ClinVar | Phelan McDermid syndrome* | Unknown significance | Not provided | 27.8 | 1.48 |
| Q317R Ank | ClinVar | Not provided | Unknown significance | Not provided | 24.48 | 0.53 |
| T337S Ank | ClinVar | 22q13 deletion syndrome* | Unknown significance | *De novo* | 52.02 | 0.11 |

\* several entries in ClinVar are listed as „Phelan-McDermid syndrome“ or under the alternative name „22q13 deletion syndrome“, though there is no deletion involved but a missense mutation, as indicated.

For biochemical analyses of Shank3 interactions, mutations were introduced into an expression construct coding for the SPN and Ank domains (residues 1-339 of Shank3) in fusion with mRFP. WT and mutant constructs, or a control vector coding for mRFP alone, were coexpressed in 293T cells with Shank3 interaction partners of interest. Cells were lyzed, and mRFP-tagged proteins were immunoprecipitated using the RFP-trap matrix. Input and precipitate samples were analyzed for the presence of Shank3 and its interaction partners by Western blotting.

In a first set of experiments, we analyzed binding to the small G-proteins Rap1 (expressed as a GFP-tagged, constitutively active G12V mutant) and H-Ras (as HA-tagged, active G12V variant). Both have been shown to interact with the SPN domain of Shank proteins in their active, GTP bound forms (17). Here, both Rap1a and HRas were efficiently coimmunoprecipitated from lysates of cells expressing WT and all variant forms of the Shank3 N-terminus. A quantitative analysis showed that there were no significant differences between mutants, suggesting that none of the mutations analyzed here has any effect on small G-protein binding (Fig. 2).

**Figure 2.**
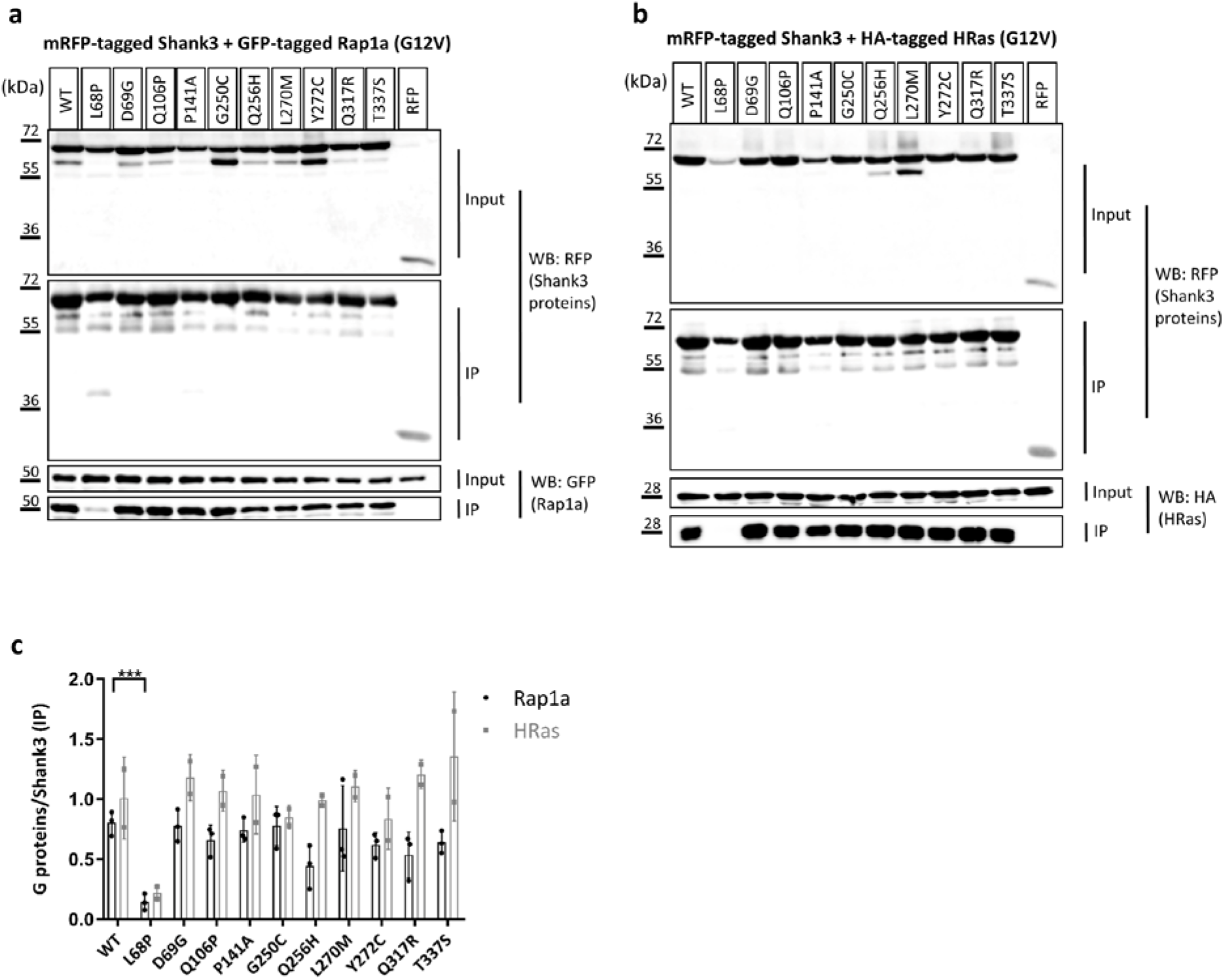
RFP-tagged WT and mutant variants of the Shank3 N-terminus (SPN + Ank domains), or mRFP alone, were coexpressed in 293T cells with active (G12V-mutant) variants of the small G-proteins Rap1a (GFP-tagged; **a**) or HRas (HA-tagged, **b**). After cell lysis, RFP-tagged proteins were immunoprecipitated using the mRFP-trap matrix. Input and precipitate samples were analysed by Western blotting using epitope-specific antibodies. **c**. Quantitative analysis of the data shown in **a** and **b**. Signal intensities in IP samples for G-proteins were divided by IP signals for mRFP-Shank3 variants. No significant differences between mutants and WT Shank3 were detected for Rap1a (n=3) and HRas (n=2), with the exception of the L68P mutant which we have shown before to unfold the SPN domain (17). ***, significantly different from WT, p<0.001; data from three (Rap1a) or two (HRas) independent experiments; ANOVA, followed by Dunnett’s multiple comparisons test.

In the next step we determined binding to δ-catenin, which we have recently identified as interaction partner of the N-terminus of Shank3 (20). Binding to δ-catenin was not affected by the D69G mutation in the SPN domain, but was significantly reduced by several mutations in the Ank repeats (e.g. L270M) (Fig. 3), consistent with our previous observation that the Ank repeats of Shank3 constitute the binding interface for δ-catenin (20).

**Figure 3.**
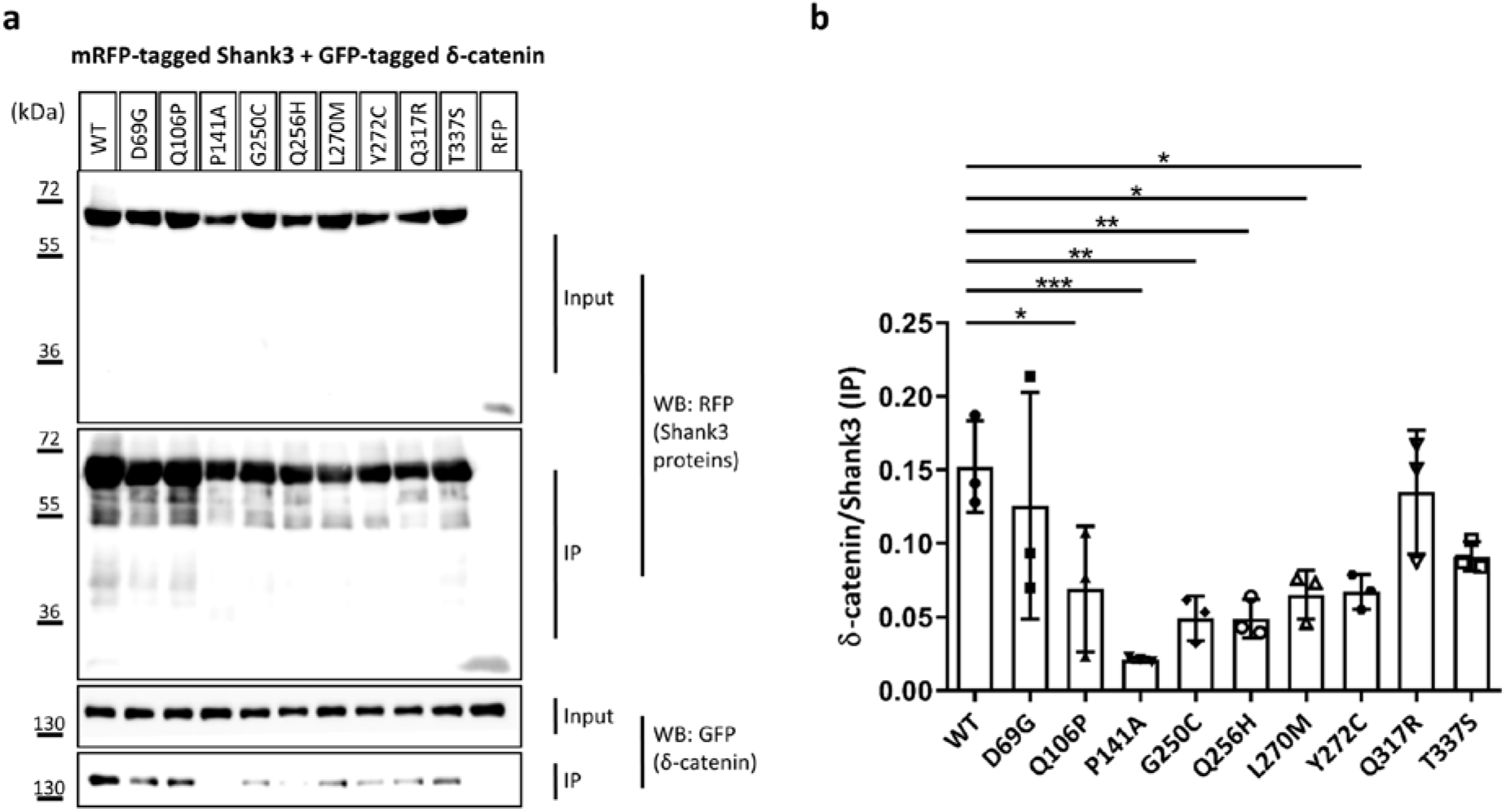
**a**. RFP-tagged WT and mutant variants of the Shank3 N-terminus (SPN + Ank domains), or mRFP alone, were coexpressed in 293T cells with GFP-tagged δ-catenin. After cell lysis, RFP-tagged proteins were immunoprecipitated using the mRFP-trap matrix. Input and precipitate samples were analysed by Western blotting using mRFP- (upper panels) and GFP-specific antibodies (lower panels). **b**. Quantitative analysis. Signal intensities in IP samples for δ-catenin were divided by IP signals for mRFP-Shank3 variants. *, **, ***, significantly different from WT, p<0.05, 0.01, p<0.001, respectively; data from three independent experiments; ANOVA, followed by Dunnett’s multiple comparisons test.

Similarly, previous work had shown that the cytoskeletal protein α-Fodrin binds to the Ank repeats of Shank proteins (19). However, our structural and biochemical studies had shown that the SPN domain, through its intramolecular interaction, blocks access of α-Fodrin to the Ank repeats (17, 22). Our current experiments supported this idea as binding of α-Fodrin to the WT N-terminus of Shank3 appeared to be rather weak. However, one of the mutations tested, P141A, strongly increased the binding of α-Fodrin to Shank3 (Fig. 4). This indicated that this mutation might open up the tight interaction between SPN and Ank repeats, allowing for better access of α-Fodrin to its binding site on the Ank repeats. In agreement, we observed that a residue at this position exhibits very low solvent accessibility (Table 1).

**Figure 4.**
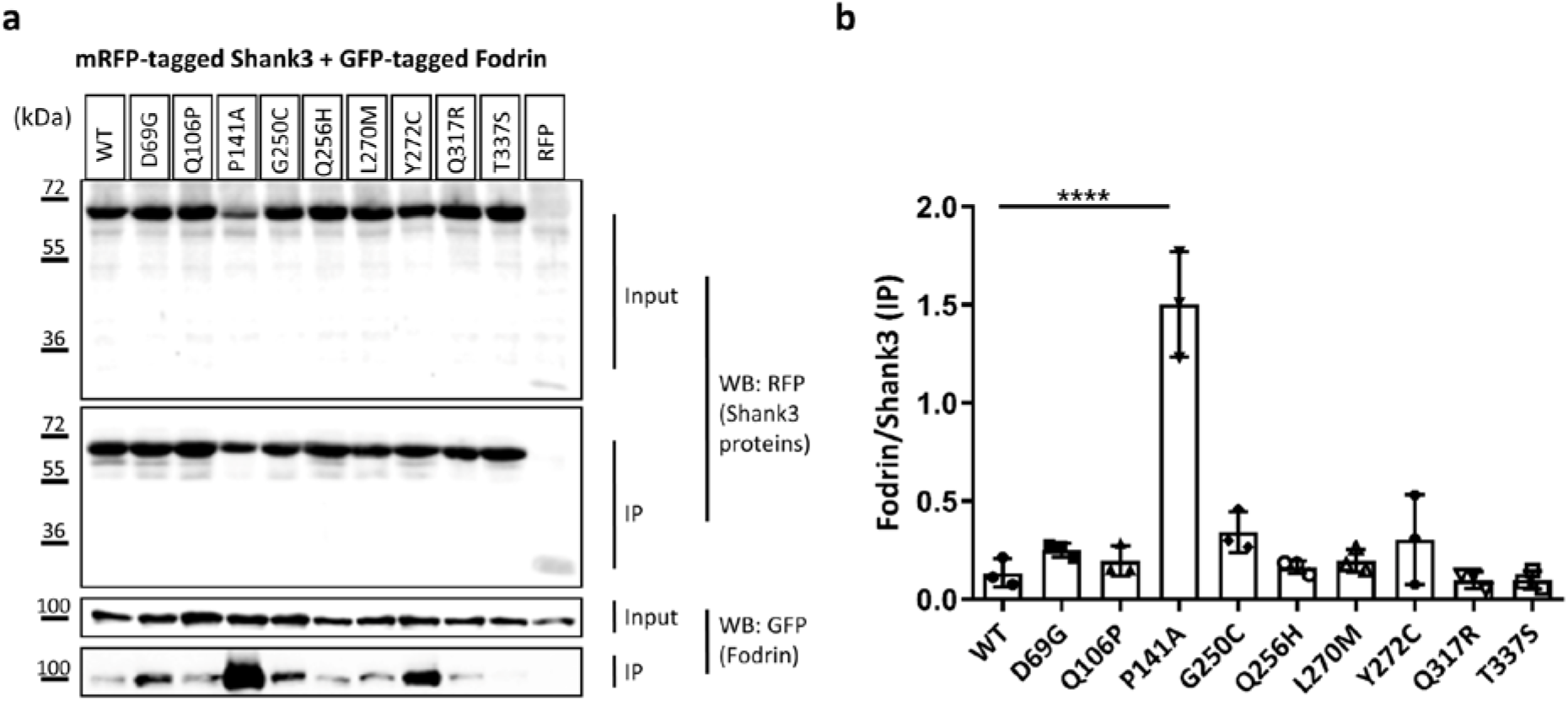
**a**. RFP-tagged WT and mutant variants of the Shank3 N-terminus (SPN + Ank domains), or mRFP alone, were coexpressed in 293T cells with a GFP-tagged C-terminal fragment of α-Fodrin (19, 22). After cell lysis, RFP-tagged proteins were immunoprecipitated using the mRFP-trap matrix. Input and precipitate samples were analysed by Western blotting using mRFP- (upper panels) and GFP-specific antibodies (lower panels). **b**. Quantitative analysis. Signal intensities in IP samples for α-Fodrin were divided by IP signals for mRFP-Shank3 variants. ****, significantly different from WT, p<0.0001; data from three independent experiments; ANOVA, followed by Dunnett’s multiple comparisons test.

This result prompted us to ask whether mutations might affect the intramolecular interaction between SPN and Ank domains. As a first approach for this, we coexpressed the GFP-tagged SPN domain with the full length N-terminus (SPN+Ank), reasoning that any mutation that would open up the SPN-Ank fold would allow for easier access of the exogenous SPN domain to the Ank repeats. Here we used the N52R mutation as a positive control. The change of Asn52 to Arg in the SPN domain has been designed based on the 3D-structure, with the intention to weaken the affinity of the SPN to the Ank domain. Indeed we have shown before that it truly opens up the SPN-Ank interaction (30). Upon coexpression of WT and mutant forms of the complete N-terminus with the GFP-tagged SPN domain, only the N52R positive control induced a significant increase in binding of the SPN domain to the N-terminus (Fig. 5). However, we realized that an opening of the SPN-Ank fold which would be induced by a loss of affinity for the SPN domain in the Ank repeats might not be detected in this experiment. To overcome this problem, we analyzed directly the interaction between Ank repeats and the SPN domain by generating an mRFP-tagged construct coding for the Ank repeats alone, which was coexpressed together with the GFP-tagged SPN domain. Here, we observed that the P141A mutation induced a strong loss of interaction with the SPN domain (Fig. 5). This observation clearly supported the idea that this mutation interferes with the SPN/Ank interaction and thereby allows for improved binding of α-Fodrin to the Ank repeats.

**Figure 5.**
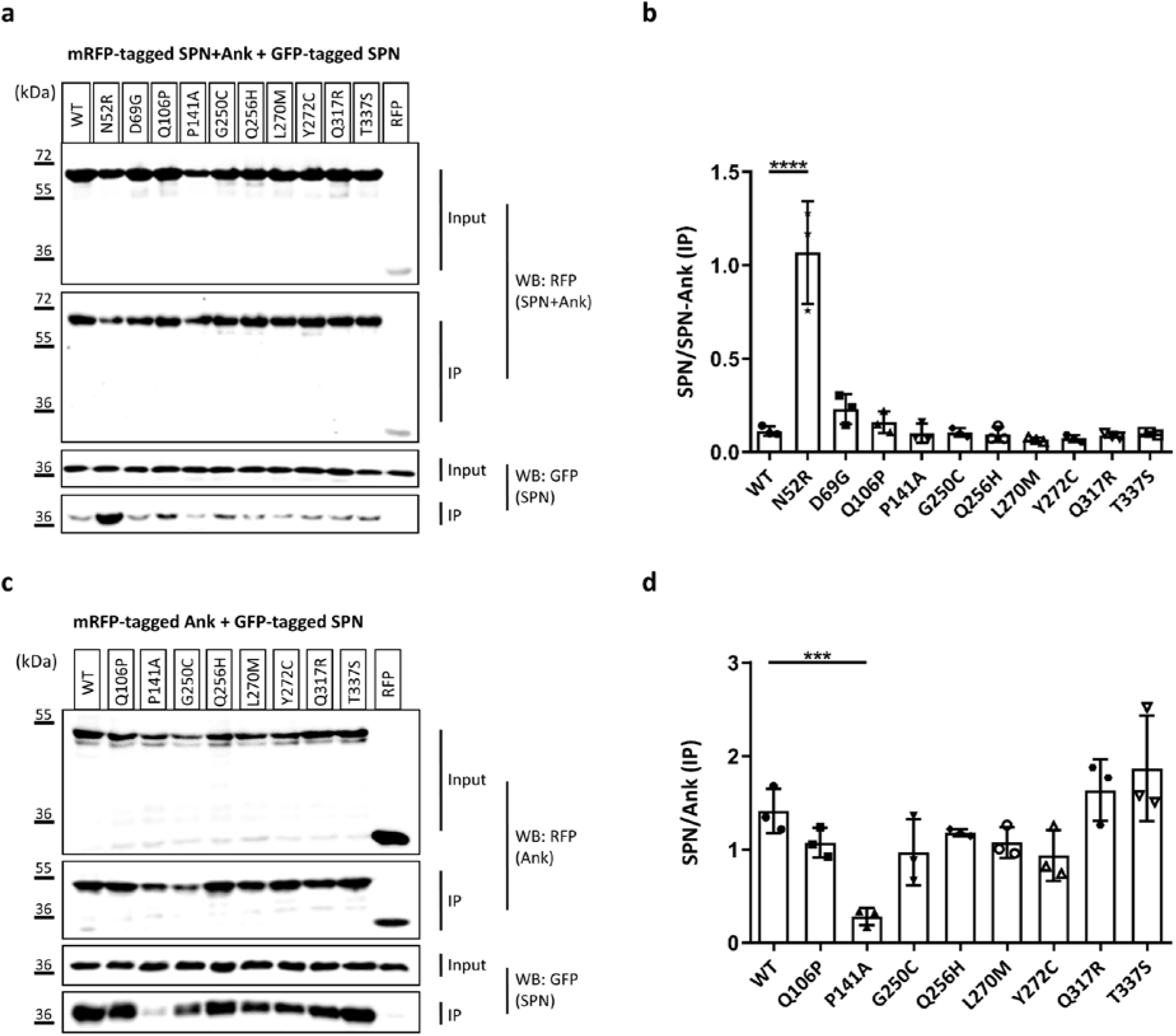
**a**. RFP-tagged WT and mutant variants of the Shank3 N-terminus (SPN + Ank domains), or mRFP alone, were coexpressed in 293T cells with the GFP-tagged SPN domain of Shank3. The N52R mutant, which was designed based on the 3D structure of Shank3 to disrupt the SPN-Ank contact, was included as a positive control here. After cell lysis, RFP-tagged proteins were immunoprecipitated using the mRFP-trap matrix. Input and precipitate samples were analysed by Western blotting using mRFP- (upper panels) and GFP-specific antibodies (lower panels). **b**. Quantitative analysis. Signal intensities in IP samples for GFP-SPN were divided by IP signals for mRFP-Shank3 variants. ****, significantly different from WT, p<0.0001; ANOVA, followed by Dunnett’s multiple comparisons test. **c**. The experiment was repeated as in **a**, however using a Shank3 construct coding for Ank repeats only. **d**. Quantitative analysis performed as in **b**. ***, significantly different from WT, p<0.001; data from three independent experiments; ANOVA, followed by Dunnett’s multiple comparisons test.

We used several biophysical methods to verify that the closed conformation of the SPN-Ank tandem is indeed opened up by some mutations found in patients. In the 3D structure of the Shank3 N-terminal domains, the N- and C-termini of the protein are located rather close to each other, indicating that FRET might be an ideal technique to monitor conformational changes. We generated FRET reporter constructs by fusing EGFP to the N-terminus, and mCherry to the C-terminus of the Shank3 fragment (residues 1-339, as before). After transfection in 293T cells, the WT fragment indeed showed high FRET efficiency, supporting the notion that both fluorophores are in close proximity to each other (Fig. 6a). We analyzed the P141A mutation, together with two positive controls for which we were confident that they would disrupt the intramolecular contact between SPN and Ank domains: N52R (see above, and(30)), and L68P, a patient mutation for which we have shown that it disrupts the folding of the SPN domain (17, 22, 31). We also included, in a separate experiment, a mutant which is not involved in the interface between SPN and Ank: R12C. We observed a significant decrease in FRET efficiency for P141A, similar to N52R and L68P, confirming that the P141A mutation opens up the SPN-Ank fold (Fig. 6a). In contrast, R12C had no effect, in agreement with its location in the Ras binding site, pointing away from the SPN-Ank interface (Fig. 6b).

**Figure 6.**
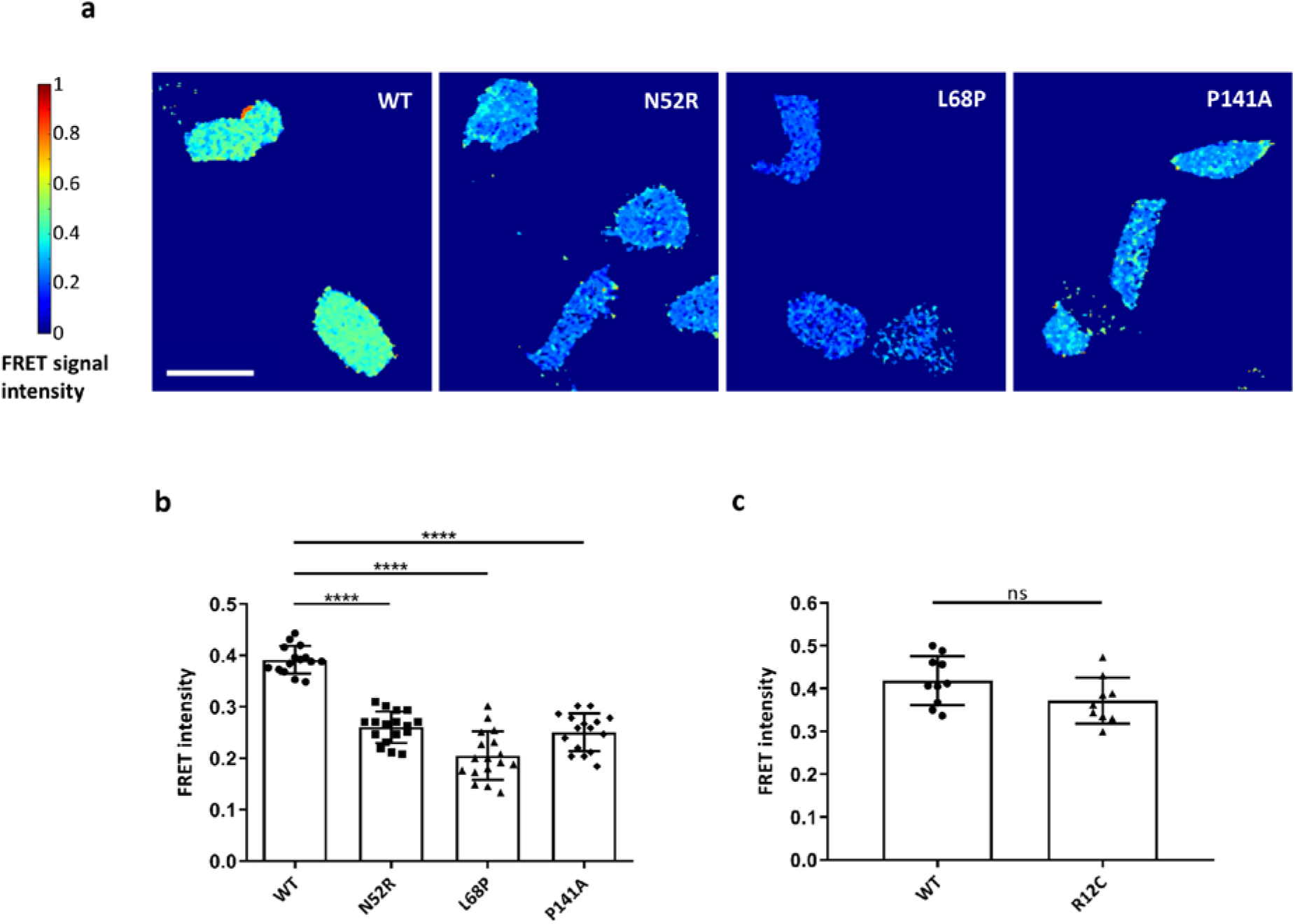
**a**. 293T cells transfected with Shank3 N-terminal FRET constructs coding for a fusion of N-terminal GFP, followed by WT or mutant Shank3 (residues 1-339, SPN+Ank), and C-terminal mCherry, were subjected to live cell imaging using a FRET imaging program (scale bar 50 μm). The FRET intensity was compared after 1 min. The color-coded FRET signal intensity (left) represents a low FRET signal (FRET=0) with the dark blue color and a very high FRET signal FRET=1 with the dark red color. **b**. Quantitative evaluation of cells shown in **a**. **c**. Repeat of the experiments shown in **a**, **b** with the R12C mutant (which is not in the interface between SPN and Ank domains). ****, p<0.0001; analysis of n=15-17 cells (**a, b**) or n=9-10 cells (**c**) from 3 independent experiments.

In a second approach, we performed differential scanning fluorimetry, which measures the thermal unfolding of the protein. Protein samples were subjected to increasing temperatures, causing an unfolding event in the protein. The SYPRO Orange dye binds to hydrophobic regions exposed as the protein unfolds, and changes in the fluorescence are monitored as the temperature increases to generate the melting curves. The Tm calculated is the temperature at which the concentration of the folded state of the protein equals the concentration of the unfolded state of the protein. The P141A protein showed a decrease in Tm of six degrees, compared to that of the WT 1-348 Shank3, indicating a substantial reduction in stability of the mutant protein compared to WT at 0.5 M NaCl conditions (Fig. 7).

**Figure 7.**
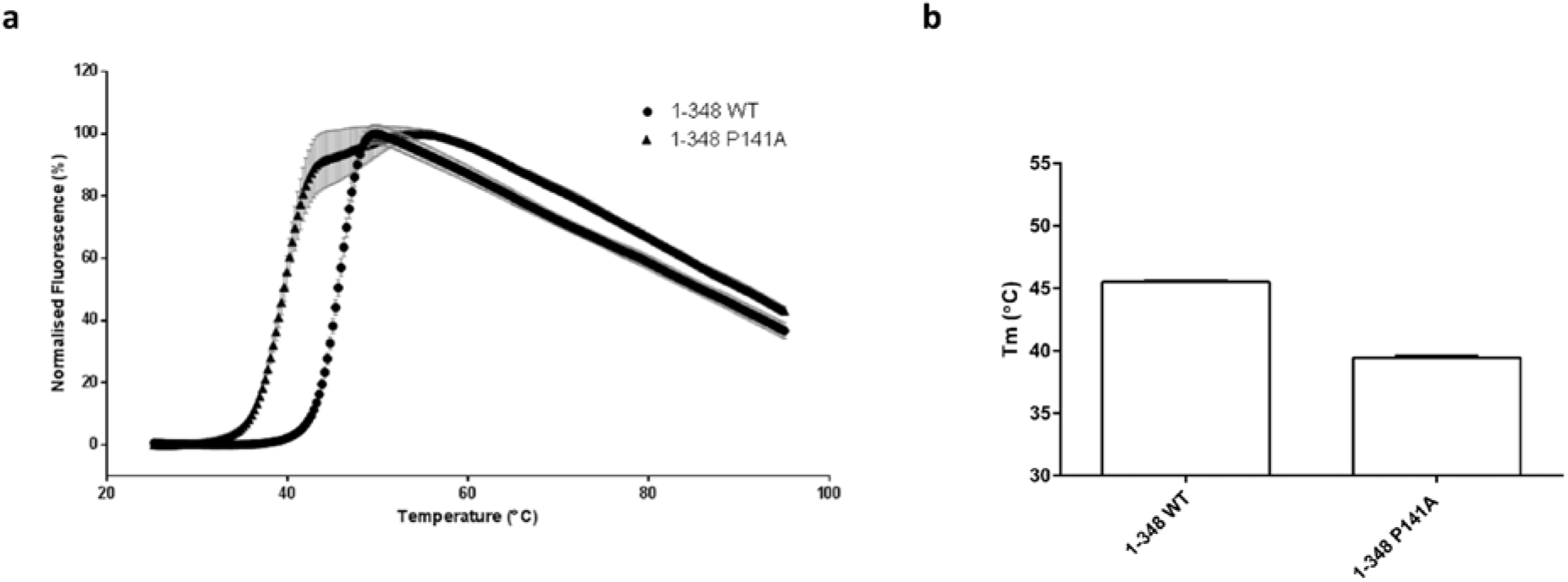
DSF analysis of stability of Shank3 constructs (1-348 WT and 1-348 P141A) in 0.5 M NaCl, 20 mM Tris, 2 mM DTT, pH 7.5. Fluorescence based on SYPRO-Orange dye. Mean normalised data plotted showing melt curves (**a**) and Tm values calculated from V50 Boltzman calculation from mean fluorescence data (**b**); n=3.

Finally, we turned our attention to the α-subunit of the calcium/calmodulin dependent kinase II (αCaMKII). In contrast to the aforementioned interaction partners, this protein, in its autoinhibited inactive form, does not specifically interact with one of the two clearly defined domains (SPN and Ank) but rather binds to the linker connecting the two domains and to the SPN domain (23). Upon coexpression and coprecipitation of Shank3 fragments with this protein, we observed strong and specific binding (Fig. 8). This was reduced by the L68P mutation (unfolding the SPN domain, thus supporting the relevance of the SPN surface for interaction with αCaMKII). In addition, both Q106P and P141A showed significantly impaired binding to the αCaMKII. For Q106P, this may be explained by the close proximity of Q106 to the linker region and the actual αCaMKII binding site on Shank3. In the original publication on this interaction (23), mutation of Ile102, which is in close proximity to Q106, also interfered strongly with this interaction. On the other hand, P141 is not localized in the linker region but is part of the interface between Ank and SPN domains. Our previous data (see Figs. 5 and 6) suggest that the primary effect of P141A is to disrupt the SPN-Ank intramolecular contact. We speculated that P141A affects αCaMKII binding by pushing SPN and Ank away from each other, thereby altering the conformation of the linker region which appears to act as a hinge between SPN and Ank repeats. To validate this hypothesis, we assessed the only other mutation which affects the SPN-Ank interface, namely the N52R mutation, for its effect on αCaMKII binding. Indeed, we observed that this mutation, which also alters a residue in the interface between Ank and SPN domains, destroyed binding of the αCaKKII.

**Figure 8.**
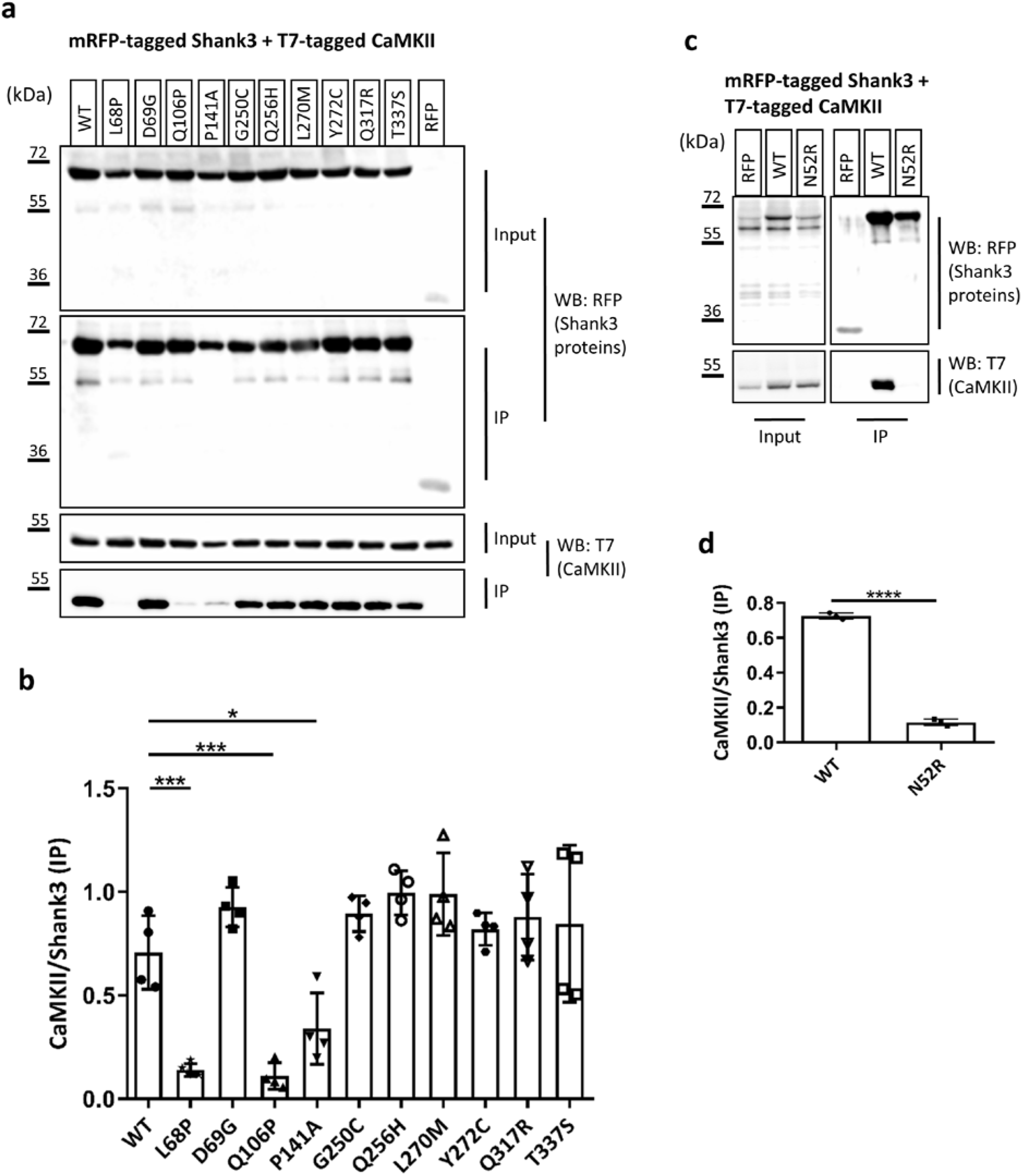
**a**. RFP-tagged WT and mutant variants of the Shank3 N-terminus (SPN + Ank domains), or mRFP alone, were coexpressed in 293T cells with T7-tagged αCaMKII. After cell lysis, RFP-tagged proteins were immunoprecipitated using the mRFP-trap matrix. Input and precipitate samples were analysed by Western blotting using mRFP- (upper panels) and T7-specific antibodies (lower panels). **b**. Quantitative analysis. Signal intensities in IP samples for T7-αCaMKII were divided by IP signals for mRFP-Shank3 variants. *, **, ***, significantly different from WT, p<0.05, 0.01, p<0.001, respectively; data from four independent experiments; ANOVA, followed by Dunnett’s multiple comparisons test. **c**, **d**. The assay as in **a**, **b** was repeated for the N52R mutation in Shank3. ****, significantly different from WT, p<0.0001; data from three independent experiments; t-Test.

In the next step we analyzed the potential relevance of our findings for synaptic function; here we focused on the P141A mutant as it had the strongest effect on protein interactions. GFP-tagged full-length Shank3 (WT and P141A mutant) was expressed in primary cultured hippocampal neurons, and transfected cells were stained and analyzed for the distribution of MAP2 (as a dendritic marker), PSD-95 (as a postsynaptic marker) and GFP-Shank3. Both WT and mutant protein were efficiently targeted to postsynaptic sites, as the GFP-Shank3 signal colocalized extensively with the PSD-95; also, we observed the Shank3- and PSD-95 positive, presumably postsynaptic clusters mostly in some distance from the (MAP2-labelled) dendritic shaft, indicating a localization on dendritic spines. However, a clear difference between WT and mutant was observed with respect to the number of these postsynaptic clusters, as we found this to be significantly reduced for neurons expressing the P141A mutant (Fig. 9).

**Figure 9.**
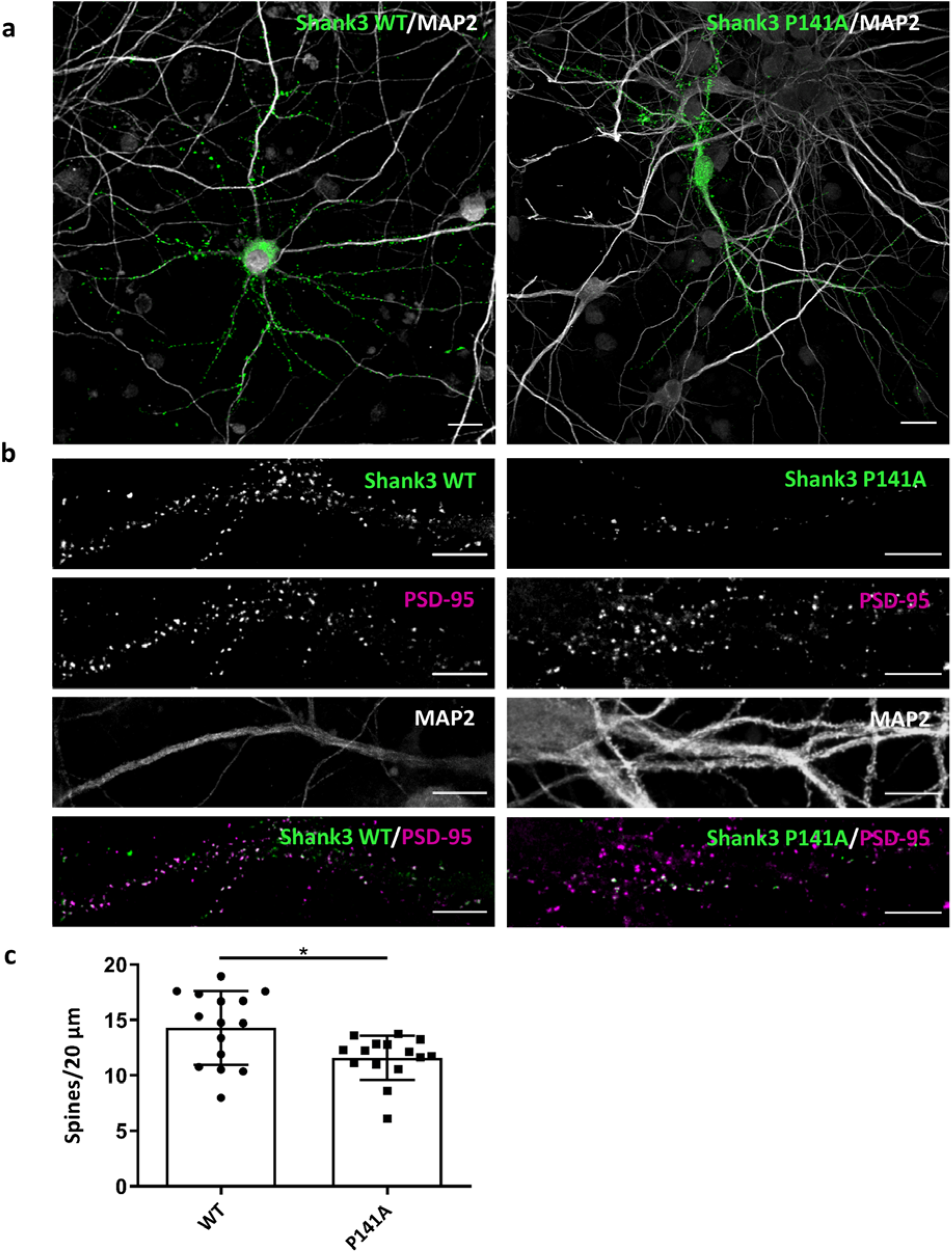
Primary hippocampal neurons expressing GFP-Shank3 (WT or P141A mutant) were fixed and stained for the postsynaptic marker PSD-95 and the dendritic marker MAP2 (gray). The expressed Shank3 variants are detected using the GFP fluorescence. **a**. Overview images of neurons (scale bar 20 μm). **b**. Magnified dendritic areas (scale bar 5 μm). GFP-Shank3 colocalizes extensively with PSD-95 in presumably postsynaptic clusters, which are found in a small distance from the (MAP2-positive) main dendrite and are therefore classified as dendritic spines. **c**. Quantitative evaluation of the number of dendritic spines per 20 μm dendrite. *, significantly different from WT, p<0.05; analysis of 45 dendrites of n=15 neurons from three independent experiments; t-Test.

## Discussion

Mutations in *SHANK3* have been mostly associated with autism and intellectual disability syndromes (16); we report here a family with the p.L270M variant, where carriers of this variant exhibit a somewhat divergent phenotypic spectrum. This includes learning difficulties (which is common in cases with *SHANK3* mutations), but also an oppositional behavioral disorder and ADHD–like behavior. ADHD has so far not been associated with alterations in any of the *SHANK* genes. In addition, we observed obesity in all carriers of this variant, a feature which has so far not been linked to alterations in *SHANK3.* The fact that this variant segregates with the phenotype suggests that it is causative; however, to prove pathogenicity we performed a number of functional assays. For this, we included further variants in the N-terminal part of SHANK3 which had so far not been functionally studied, including several mutations which had been deposited in the ClinVar database with only limited phenotypical description.

We aimed for a thorough biochemical analysis of the effects of mutations affecting the N-terminal parts of the Shank3 protein. For this we used a standardized assay of coexpression/co-immunoprecipitation of various known interaction partners of Shank3, with an mRFP-tagged construct which contains the SPN and Ank repeat domains. We used this very “reduced” construct because we wanted to study the isolated effects of N-terminal mutations. It should be kept in mind that some interaction partners make more than one contact to the full length Shank3 protein; thus, the α-subunit of CaMKII which was studied here binds to the linker between SPN domain and Ank repeats in its inactive, non-phosphorylated form (23). In addition, it binds to a short sequence segment within the proline rich region in its active form (32). Thus, any isolated effect of mutations such as the Q106P mutation might have been lost in the context of the full-length protein.

The effects we observed may be divided into two categories (which are not mutually exclusive): several mutations had a negative impact on the interaction with δ-catenin. As we had identified the Ank repeats of Shank3 as the domain binding to δ-catenin, it is conceivable that the mutations alter the interface between δ-catenin and Shank3 and thereby reduce binding. As a reduction in δ-catenin interaction was the only effect observed with the L270M mutation, it may be argued that loss of δ-catenin binding only may be associated with the somewhat milder phenotype exerted by this mutation. However, such a conclusion would require analysis of more patients/variants.

Another set of mutations interfered with the closed conformation between SPN domain and Ank repeats, and/or the binding of the α-subunit of the CaMKII to the linker between both domains. One intriguing finding we made here is that these two aspects are intricately linked to each other. Thus we observed that, upon disruption of the SPN-Ank contact using N52R or P141A mutations, we also lose binding of αCaMKII (see model in Fig. 10). Loss of αCaMKII was also observed with the R12C and L68P mutations which have been more widely studied. R12, similar to Q106 analyzed here, appears to be in direct contact with αCaMKII (23), whereas the L68P mutation leads to a local unfolding of the SPN domain ((17, 22, 31)). As loss of αCaMKII binding appears to be a more common result of *SHANK3* missense mutations, one might ask what the physiological relevance of this interaction may be. Currently, we speculate that Shank3 acts as a negative regulator of the αCaMKII signalling pathway which is prominently involved in synaptic plasticity. One prediction here would be that Shank3 deficiency (or the presence of mutants R12C, L68P or P141A) would lead to a higher basal activity of αCaMKII due to loss of the inhibiting effect of the Shank3 N-terminus.

**Figure 10.**
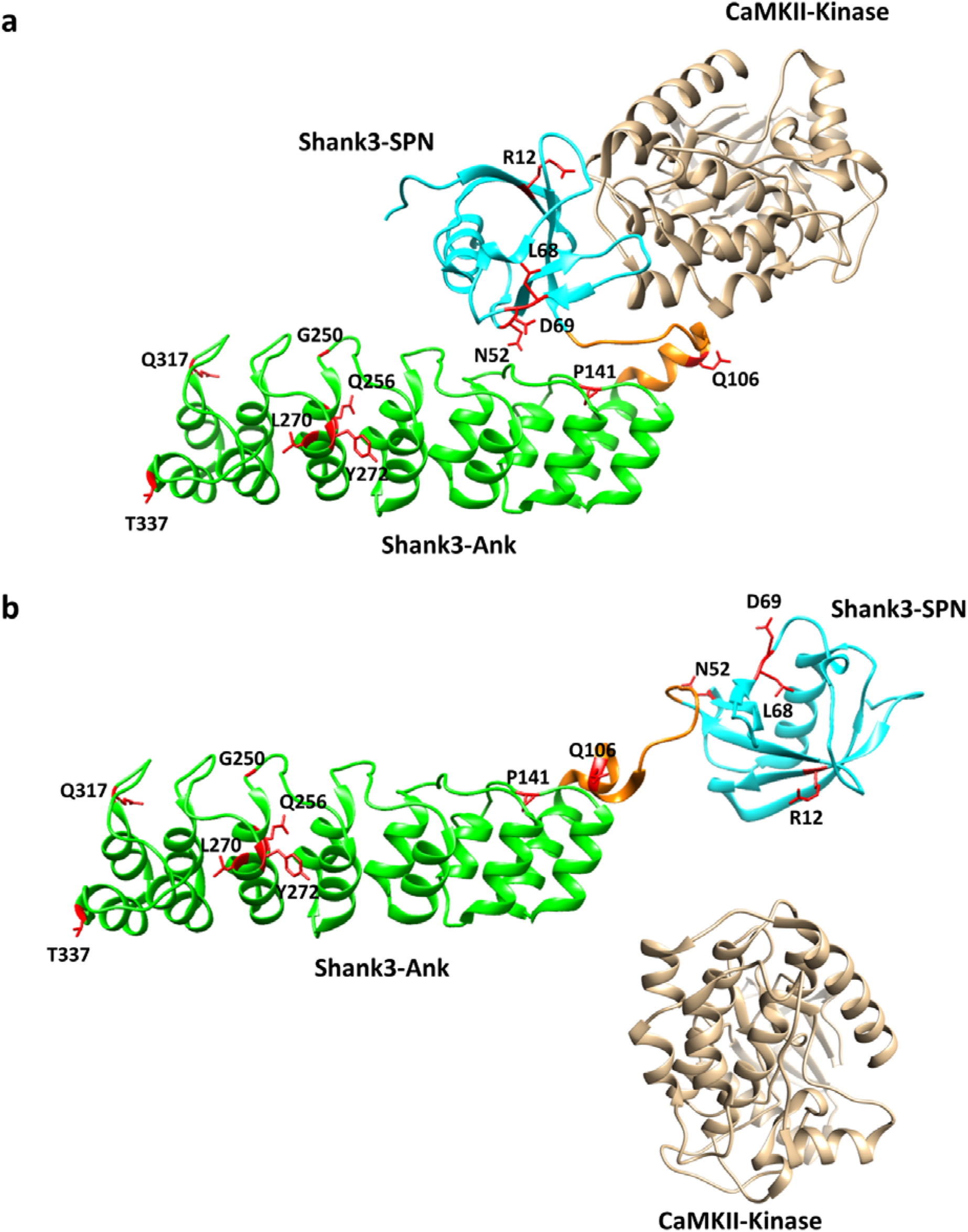
Model of open/closed transition of the N-terminal domains of the Shank3 proteins. **a**. In the closed conformation, a shallow cavity is formed between the SPN domain, the linker between SPN and Ank, and the N-terminal end of the helix beginning with Q106. Bound αCaMKII covers a large part of this interface through interactions with both the SPN and the linker sequence, including and up to Q106. **b**. Upon disruption of the SPN-Ank interaction, the geometry of the αCaMKII binding site is altered and αCaMKII cannot bind anymore. Model based on the crystal structure of the Shank3 N-terminus (5G4X)(17) and models of the Shank3/ αCaMKII complex shown in ref. (23).

Importantly, it appears that the p.P141A variant of *SHANK3* has the most dramatic effect on the function of the protein, as it disrupts the intramolecular interaction between SPN and Ank, and interferes with αCaMKII and δ-catenin binding. Our biophysical analyses, using FRET and DSF techniques, support this conclusion. They also show that the separation of domains destabilizes the protein, as we recorded a lower melting point of the Shank3 N-terminus in the DSF assay. Importantly, the P141A variant reduces the capacity of Shank3 to support the formation of postsynaptic clusters on dendritic spines, as the number of these clusters is reduced upon expression in cultured hippocampal neurons. Thus, P141A and L270M may stand for two opposite ends of the mutational spectrum in this part of the Shank3 protein. P141A is associated with multiple disruptions of N-terminal interactions; this mutation was not inherited but occurred *de novo* in the patient and causes a rather severe phenotype with intellectual disability and autism. On the other hand, L270M affected only one of the interactions tested here; this variant was inherited and causes a comparatively mild phenotype involving ADHD and learning difficulties.

## Abbreviations

Abi-1: Abl interactor 1
ADHD: Attention deficit hyperactivity disorder
Ank: ankyrin repeats
ANOVA: analysis of variance
ASD: autism spectrum disorder
CaMKII: Ca2+/calmodulin-dependent protein kinase II
cDNA: complementary deoxyribonucleic acid
chk: chicken
DAPI: 4’,6-Diamidino-2-phenylindole
DIV: Days *in vitro*
DSF: differential scanning fluorimetry
DTT: Dithiothreitol
EDTA: Ethylenediaminetetraacetic acid
EGFP: enhanced green fluorescent protein
FRET: Förster energy resonance trasfer
GFP: green fluorescent protein
GKAP: guanylate kinase-associated protein
GTP: guanosine triphosphate
HA: Human influenza hemagglutinin
HBS: Hepes buffered salt solution
HBSS: Hank’s balanced salt solution
HEK: Human embryonic kidney
HRP: horseradish peroxidase
ICC: immunocytochemistry
ID: intellectual disability
IgG: immunoglobulin G
IP: immunoprecipitation
IRSp53: insulin receptor substrate 53
MAP2: microtubule-associated protein 2
mRFP: monomeric red fluorescent protein
ns: not significant
PAGE: polyacrylamide gel electrophoresis
PBS: phosphate-buffered saline
PCR: polymerase chain reaction
PDZ: PSD-95/DLG/ZO1
PSD: post-synaptic density
PSD-95: postsynaptic density protein 95
SAM: sterile alpha motif
SAPAP: SAP90/PSD-95-associated protein
SD: standard deviation
SDS: sodium dodecyl sulfate
Shank: SH3 and multiple ankyrin repeats
SPN: Shank3/ProSAP N-terminal
SUMO: Small Ubiquitin-like Modifier
TBS-T: tris-buffered saline-Tween 20
Ubl: ubiquitin like
WES: Whole exom sequencing
WB: Western blot
WT: wild type

## Acknowledgements

The authors thank Hans-Hinrich Hönck (UKE Hamburg) for excellent technical assistance. The work was supported by grants from Deutscher Akademischer Austauschdienst (DAAD; to F.H.-N.) and Deutsche Forschungsgemeinschaft (DFG; KR 1321/9-1; to H.-J.K). C.F. was supported by the region of Southern Denmark.

