## Supplemental data for "Mutations affecting the N-terminal domains of SHANK3 point to different pathomechanisms in neurodevelopmental disorders"

**Patient data.**

**Individual III:1.** The index patient is a boy aged 12 years born as the first child of non-consanguineous parents. He was born at gestational age 36+0 weeks with a birth weight of 2566 g, length 50 cm, head circumference 32 cm, and Apgar score 10/1, 10/5. He had from early on a poor sleep with several nightly awakenings. He had language delay. He was referred to the pediatric department at age 2½ years due to overweight. Poor balance, and limited language, concentration difficulties, and an aggressive behavior. He walked at age 1 year and had recurrent middle ear infections. Already as an infant he showed outward reacting behavior, and started in a special group in kindergarden, school is a special school. At age 6½ years he was seen with premature adrenarche. A slight hypermetropia was noted. He lives in an institution and has done that in periods since age 6 years. A WPSSI-R test at age 4 years showed a total IQ of 92. A WISC-IV-test at age 11 years showed a total IQ of 73 with an uneven profile extending from 66 for processing speed to 97 for reasoning, the low threshold for frustration was suspected to influence the test. He does not fulfill the criterias for ADHD, but has an outward reacting behavior with a tendency of being involved in fights, demand avoidance, irritability, and controllable behavior. He is, on the basis of these features, evaluated to fulfill the criterias for Oppositional Defiant Disorder, ODD. He needs structure and a predictable day. Current height is 171.5 cm (+2.5 SD), weight 90.5 kg (+3.2 SD), resulting in a Body mass index (BMI) of 30.8. Head circumference at age 10 y 4 m was 55 cm (approximately +1.5 SD).

Investigation for Fragile X and methylation specific MLPA for Prader Willi Syndrome showed normal result. Chromosomal microarray at age 3 years showed a maternally inherited duplication in Xp11.14 of 140 kb with a breakpoint in *TSPAN7*, that was later evaluated to be a rare normal variant without phenotypic consequence. Duo-exome analysis revealed the maternally inherited missense variant *SHANK3* (NM_033517.1):c.808C>A, p.(Leu270Met).

**Individual II:2.** The mother of individual III:1, current age 32 years. She had severe overweight since childhood. She was a problematic child in school with outward reacting behavior. There were some learning difficulties, but she has taken elementary school exam. She had cholecystitis with cholecystectomia at age 21 years. She was diagnosed with ADHD, for which she receives medical treatment with good effect. She is also diagnosed with an emotional instable personality borderline disorder (SCIDII-interview). At age 31 years she had surgery for bilateral carpal tunnel syndrome. She is severely overweight, with height 176 cm, and weight 137 kg, resulting in a BMI of 44.

**Individual II:3.** The brother of individual II:2, current age 30 years. He has since early childhood had an outward reacting behavior, and had a problematic time in school. From 5^th^ grade, he played truant most of the time. He had difficulties in concentration, and learning disabilities especially in mathemathics, while he has performed well in language fag after medication (as an adult). He has had periodical abuse of alcohol, cannabis and narcotics since teenage years. He was diagnosed with ADHD at approximately 20 years, medical treatment had good effect, however he gained weight. He was diagnosed with Wolf-Parkinson-White. He has been sentenced to prison twice due to theft and weapon possession respectively. He has been overweight in periods. His present height is 185 cm and weight 114 kg, resulting in a BMI of 33.
